# Deregulations of miR-1 and its target Multiplexin promote dilated cardiomyopathy associated with myotonic dystrophy type 1

**DOI:** 10.1101/2022.09.06.506816

**Authors:** Anissa Souidi, Masayuki Nakamori, Monika Zmojdzian, Teresa Jagla, Yoan Renaud, Krzysztof Jagla

## Abstract

Myotonic dystrophy type 1 (DM1) is the most common muscular dystrophy. It is caused by the excessive expansion of non-coding CTG repeat, which when transcribed affect functions of RNA-binding factors. Specifically, MBNL1 is sequestered in nuclear foci while CELF1 is stabilised, with adverse effects on alternative splicing, processing and stability of a large set of muscular and cardiac transcripts. Among these effects, inefficient processing and down-regulation of muscle- and heart-specific miRNA, *miR-1*, has been reported in DM1 patients, but the impact of reduced *miR-1* on DM1 pathogenesis was unknown. Here, we used *Drosophila* DM1 models to explore *miR-1* involvement in cardiac dysfunction in DM1. We found that *miR-1* down-regulation in the heart led to dilated cardiomyopathy (DCM), a DM1-associated phenotype. We then combined *in silico* screening for *miR-1* targets with transcriptional profiling of DM1 cardiac cells to identify *miR-1* target genes with potential roles in DCM. We identified *Multiplexin* (*Mp*) as a new cardiac *miR-1* target involved in DM1. Mp and its human ortholog Col15A1 were both highly enriched in cardiac cells of DCM-developing DM1 flies and in heart samples from DM1 patients with DCM, respectively. Importantly, when overexpressed in the heart, Mp induced DCM, whereas its attenuation ameliorated the DCM phenotype in aged DM1 flies. Reduced levels of *miR-1* and consecutive up-regulation of its target *Mp/Col15A1* are thus critical in DM1-associated DCM.

## Introduction

Myotonic dystrophy type 1 (DM1) is the most common muscular dystrophy in adults, with an estimated incidence of one in 8000 births (Meola and Cardani, 2015). DM1 is a multisystem disorder, with cardiac abnormalities accounting for 30% of fatalities (Mathieu et al., 1999). Cardiac involvements in DM1 include conduction defects, supraventricular and ventricular arrhythmias (Pelargonio et al., 2002), impaired diastolic or systolic function (Hermans et al., 2012; Pelargonio et al., 2002) and dilated cardiomyopathy (DCM) (Hermans et al., 2012; Lin et al., 1989; Harold H. Nguyen et al., 1988; Papa et al., 2018; Schilling et al., 2013).

The primary cause of DM1 is gain-of-function of toxic transcripts carrying expanded non-coding CUG repeats that aggregate into foci in nuclei, sequestering RNA-binding protein Muscleblind-like 1 (MBNL1) (Fardaei et al., 2002, 2001). Reduced MBNL1 levels and concomitant stabilisation of another RNA-binding protein, CELF1 (Kuyumcu-Martinez et al., 2007), lead to deregulation of alternative splicing, with abnormal expression of embryonic splice variants in adult tissues (Lee and Cooper, 2009). For example, CELF1-dependent inclusion of fetal exon 5 in the adult isoform of *cardiac troponin T* (*cTnT*) has been associated with cardiac conduction defects in DM1 (Philips et al., 1998). Besides their roles as alternative splicing regulators, both MBNL1 and CELF1 are involved in mRNA translation (Dasgupta and Ladd, 2012; de Haro et al., 2013; Timchenko et al., 2006, 2005), de-adenylation and decay (Dasgupta and Ladd, 2012; Masuda et al., 2012; Vlasova et al., 2008), and in RNA editing (Dasgupta and Ladd, 2012). MBNL1 is specifically involved in miRNA processing (Rau et al., 2011). These diverse functions of MBNL1 and CELF1 mean that DM1 may involve deregulation of multiple pathways.

To investigate the pathophysiology and the molecular mechanisms underlying DM1, several DM1 models, both mouse (Huguet et al., 2012; Orengo et al., 2008; Wang et al., 2007) and *Drosophila* (de Haro et al., 2006; Garcia-Lopez et al., 2008; Houseley et al., 2005; Picchio et al., 2013), have been created.

Reduction of MBNL1 and stabilisation of CELF1 is thought to be involved in most DM1 phenotypes. Indeed, *Mbnl1* knockout mice develop muscle myotonia, weakness/wasting and cardiac defects including dilated cardiomyopathy and heart conduction block (Lee et al., 2013). Mice overexpressing *CELF1* in the heart show conduction abnormalities and dilated cardiomyopathy (Koshelev et al., 2010) thus confirming the contribution of MBNL1 sequestration and CELF1 upregulation to DM1 pathogenesis. Overall, the mouse models reproduced multiple DM1 features including RNA foci formation and various alternative splice defects.

We generated a series of inducible *Drosophila* DM1 lines bearing UAS-iCTG constructs with 240, 480, 600 and 960 CTGs (Picchio et al., 2013). These lines were used to model DM1 in larval somatic muscles showing not only nuclear foci formation and Mbl sequestration but also muscle hypercontraction, splitting of muscle fibres, reduced fibre size and myoblast fusion defects leading to impaired larva mobility (Picchio et al., 2013). The severity of phenotypes in these *Drosophila* models could be correlated with repeat size (Picchio et al., 2013), as also observed in DM1 patients. Finally, the overexpression of *Drosophila CELF1* ortholog *Bru-3* and attenuation of *MBNL1* counterpart *mbl* (Picchio et al., 2018; Auxerre-Plantié et al., 2019) offer further valuable models for identifying gene deregulations underlying DM1.

Among molecular mechanisms associated with DM1, deregulation of miRNAs and in particular reduced levels of evolutionarily conserved muscle- and heart-specific miRNA, *miR-1*, has been reported in DM1 patients (Rau et al., 2011) and in DM1 models including mouse (Kalsotra et al., 2014) and *Drosophila* (Fernandez-Costa et al., 2013). However, the impact of *miR-1* down-regulation on DM1-associated phenotypes has not yet been analysed.

Here, we made use of *Drosophila* DM1 models to explore *miR-1* involvement in cardiac dysfunction in DM1. We observed that *dmiR-1* level was reduced in the cardiac cells of DM1 flies and that its down-regulation in the heart led to DCM, thus suggesting that reduced *dmiR-1* levels contribute to DM1-associated DCM. Among potential *dmiR-1* regulated genes from *in silico* screening, we identified *Multiplexin* (*Mp*) / *Collagen15A1* (*Col15A1)* as a new cardiac *dmiR-1* target involved in DM1. Both Mp and Col15A1 proteins were highly enriched in cardiac cells of DCM-developing DM1 flies and in heart samples from DM1 patients with DCM, respectively. Moreover, heart-targeted overexpression of Mp was sufficient to induce DCM, whereas its attenuation ameliorated the DCM phenotype in DM1 flies. *miR-1* and its target *Mp/Col15A1* thus emerge as molecular determinants of DM1-associated DCM.

## Results

### Heart-targeted *dmiR-1* attenuation causes DCM in *Drosophila*

Reduced *miR-1* levels had previously been observed in mice developing DCM (Isserlin et al., 2014) and in cardiac samples from patients with end-stage DCM (Ikeda et al., 2007). It had also been shown that *miR-1* knockout mice display the DCM phenotype (Wei et al., 2014). However, whether *miR-1* attenuation specifically within the heart leads to DCM has not been assessed. Here we tested heart-specific knockdown (KD) of *dmiR-1* in *Drosophila* using a sponge system (Fulga et al., 2015). In the *Hand>dmiR-1KD* context, the adult fly hearts showed a larger diameter with an enlarged cardiac lumen (Fig. 1A, B). Consistent with this observation, analyses of semi-intact *Hand>dmiR-1*-*KD* heart preparations and generated M-modes (Fig.1C) confirmed enlargement of the cardiac tube and showed significantly increased diastolic and systolic heart diameters (Fig. 1D, E). We then characterized the effects of *dmiR-1* down-regulation on heart contractility by assessing percent fractional shortening (%FS), which refers to the size of the cardiac tube at the end of systole and diastole. The cardiac dilation in *Hand>dmiR-1-KD* flies was associated with a significant reduction in heart contractility (Fig. 1F). *dmiR-1* attenuation in the *Drosophila* heart thus leads to DCM.

**Figure 1.**
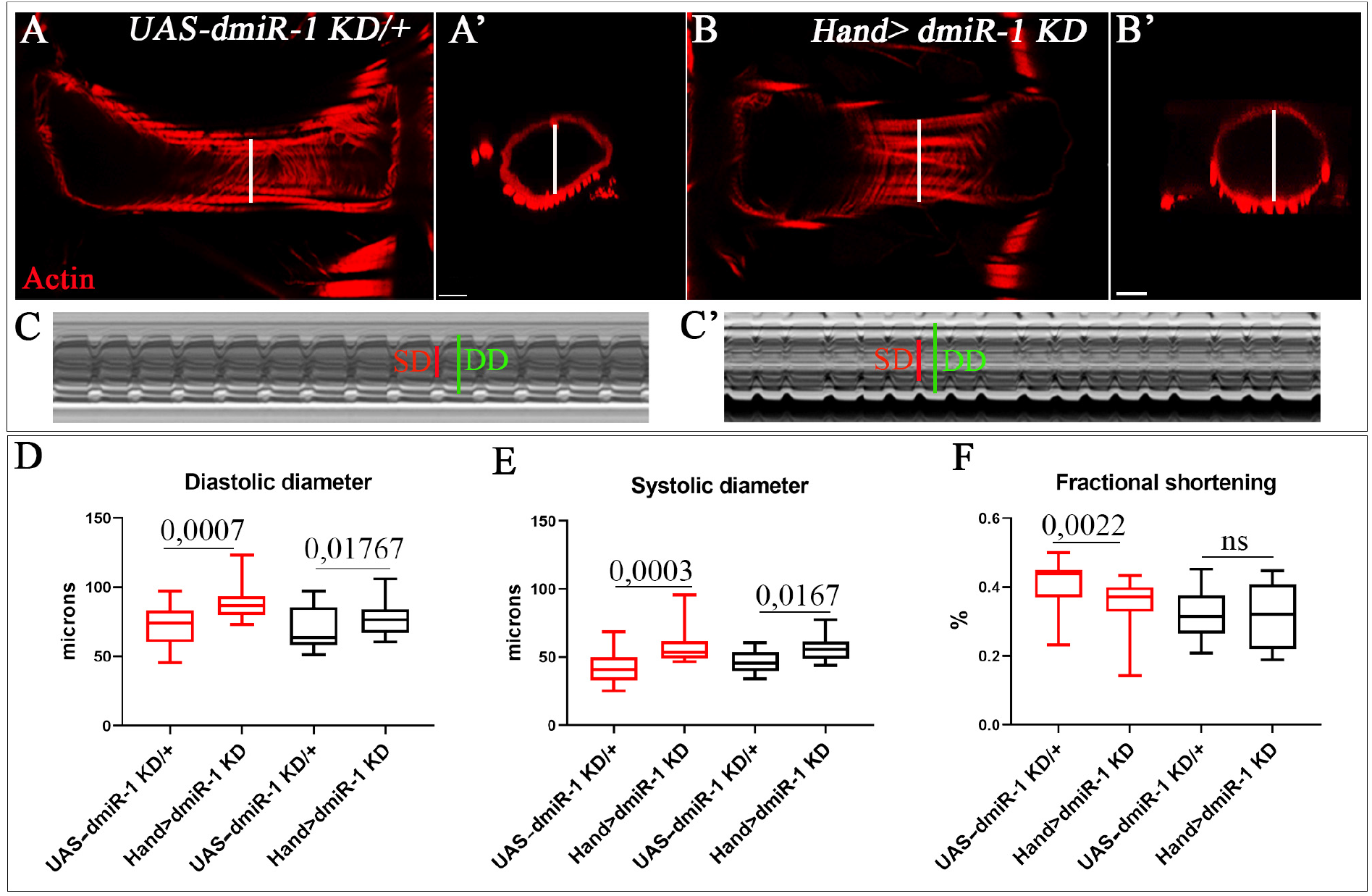
Heart-targeted *dmiR-1* attenuation causes DCM in *Drosophila*. **(A)** Control (*UAS_dmiR-1 sponge*) and **(B)** mutant (*Hand>dmiR-1 sponge*) adult hearts from one-week-old flies labelled for F-actin (red). (**A’, B’)** Cross-sections of cardiac tubes 3D-reconstructed using Imaris software. M-mode records from one-week-old control (*UAS_ dmiR-1 sponge*) **(C)** and *dmiR-1KD* (*Hand>dmiR-1 sponge*) **(C’)** flies showing increased diastolic (green) and systolic diameters (red) in the *dmiR-1KD* context **(C)**. Diastolic **(D)** and systolic **(E)** diameters and cardiac contractility analyses (percent fractional shortening) **(F)** performed using SOHA approach for control (*UAS_dmiR-1 sponge*) and *dmiR-1KD* (*Hand>sponge dmiR-1*) contexts at ages one (red) and five (black) weeks. *n* = 20 hearts. Scale bar = 20 µm.

### DM1 flies develop a DCM phenotype and show a reduced *dmiR-1* level in cardiac cells

DCM accounts for fatal cardiac involvements in DM1 patients, but the gene deregulations underlying DM1-associated DCM have not been identified. To address this issue, we first tested whether *Drosophila* DM1 models developed DCM. In three heart-specific DM1 contexts, namely (i) overexpression of 960CTG repeats, (ii) overexpression of *CELF1* orthologue *Bru3*, and (iii) attenuation of *MBNL1* orthologue *mbl*, the DCM phenotype was present in *Hand>Bru3* and *Hand>mblRNAi* models (Fig. 2A,B,C,D,E,F) but not in *Hand>960CTG* (Fig. S1A,B,C).

**Figure 2.**
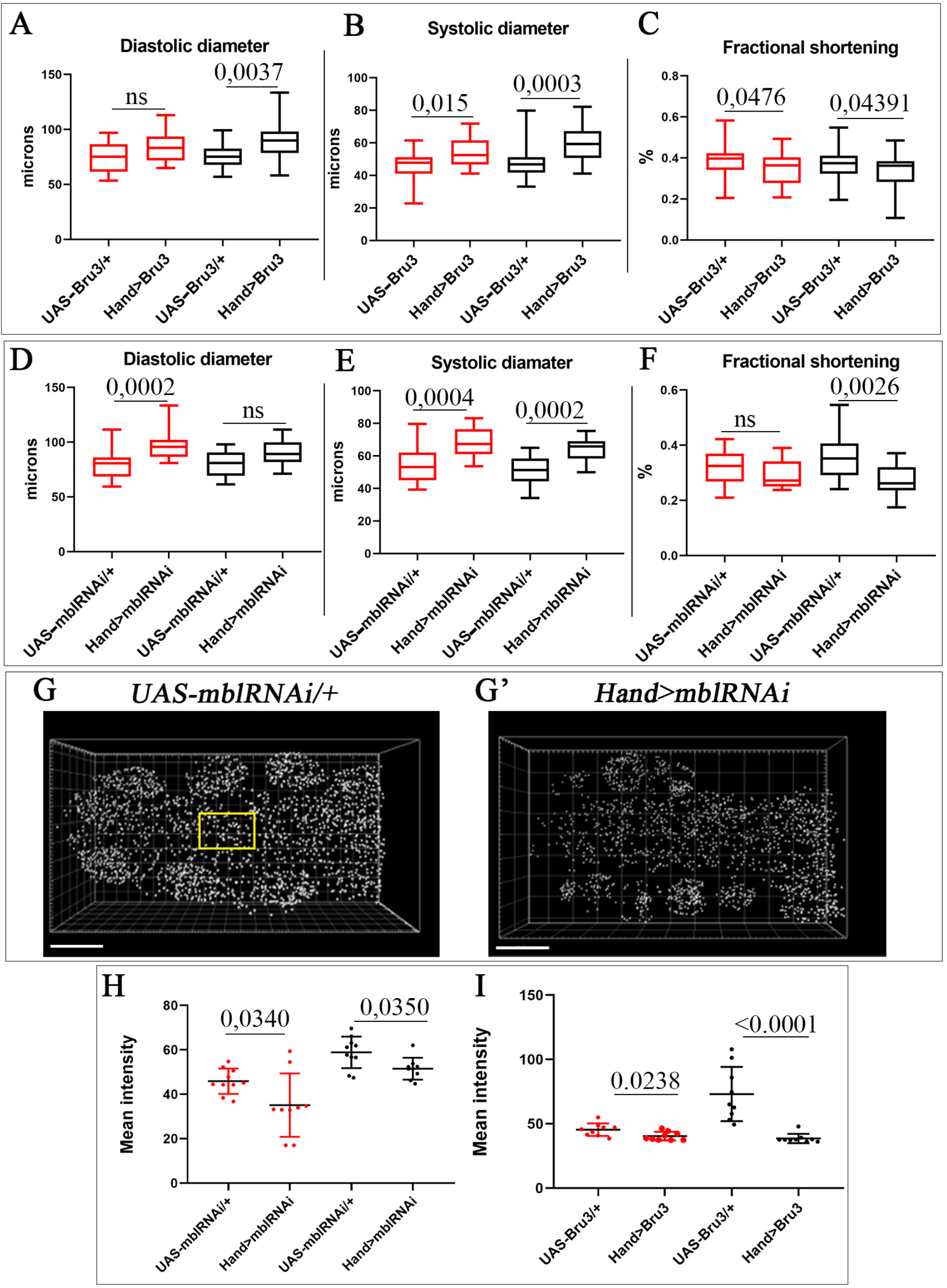
DM1 flies develop DCM phenotype with a reduced *dmiR-1* level in cardiac cells. Cardiac tube size (diastolic **(A, D)** and systolic **(B, E)** diameters) and contractility (percent fractional shortening **(C, F)**) analyses performed using SOHA approach for controls (*UAS_Bru3* and *UAS_mblRNAi*) and DM1 contexts (*Hand>Bru3* and *Hand>mblRNAi*), at ages one (red) and five (black) weeks. *n* = 20 hearts. Representative spot views generated using Imaris from *in situ* hybridisations with miRCURY LNA probe for *dmiR-1* and used for quantification of *dmiR-1* levels. Spot views of *dmiR-1* in hearts of one-week-old control (*UAS_mblRNAi*) **(G)** and DM1 flies (*Hand>mblRNAi*) **(G’)** are shown. The zoom region corresponds to area used for smFISH quantifications. **(H, I)** Scatter plot graph showing the signal intensity quantified in cardioblasts of one-(red) and five-(black) week-old flies for controls (*UAS_Bru3, UAS_mblRNAi*) and DM1 contexts (*Hand>Bru3, Hand>mblRNAi*). *n* = 9 hearts. Scale bar = 40 µm

Given that *dmiR-1* attenuation leads to DCM and that *Hand>Bru3* and *Hand>mblRNAi* DM1 models develop a DCM phenotype, we tested whether cardiac cells of *Hand>Bru3* and *Hand>mblRNAi* flies showed reduced *dmiR-1* levels. Using single molecule fluorescence *in situ* hybridisation (smFISH), we found a significantly reduced *dmiR-1* level in cardiac cells of both DCM-developing DM1 contexts (Fig. 2G,H,I and Fig. S2). *Pre-miR-1* expression was also lower in young DM1 flies (Fig. S1D), whereas hearts from aged *Hand>mblRNAi* flies showed an increased *pre-miR-1* level, most probably owing to impaired processing (Fig. S1D). DCM-developing *Drosophila* DM1 models could thus serve to test and identify genes deregulated in the DM1-associated DCM context in response to a reduced *dmiR-1* level.

### *dmiR-1* target Multiplexin is up-regulated in DCM-developing DM1 flies

To identify *dmiR-1* target genes involved in DM1-associated DCM, we first performed *in silico* screening by mapping *Drosophila*-specific *dmiR-1* seed sites annotated in miRBase (Griffiths-Jones et al., 2006; http://www.mirbase.org/) on 3’UTR regions of *Drosophila* genes up-regulated in two DCM-developing DM1 contexts (Table 1; Fig. 3A) (Auxerre-Plantié et al., 2019). We identified a set of candidate genes, of which *Multiplexin (Mp)* (Table 1, scheme in Fig. 3A).

**Table 1.**
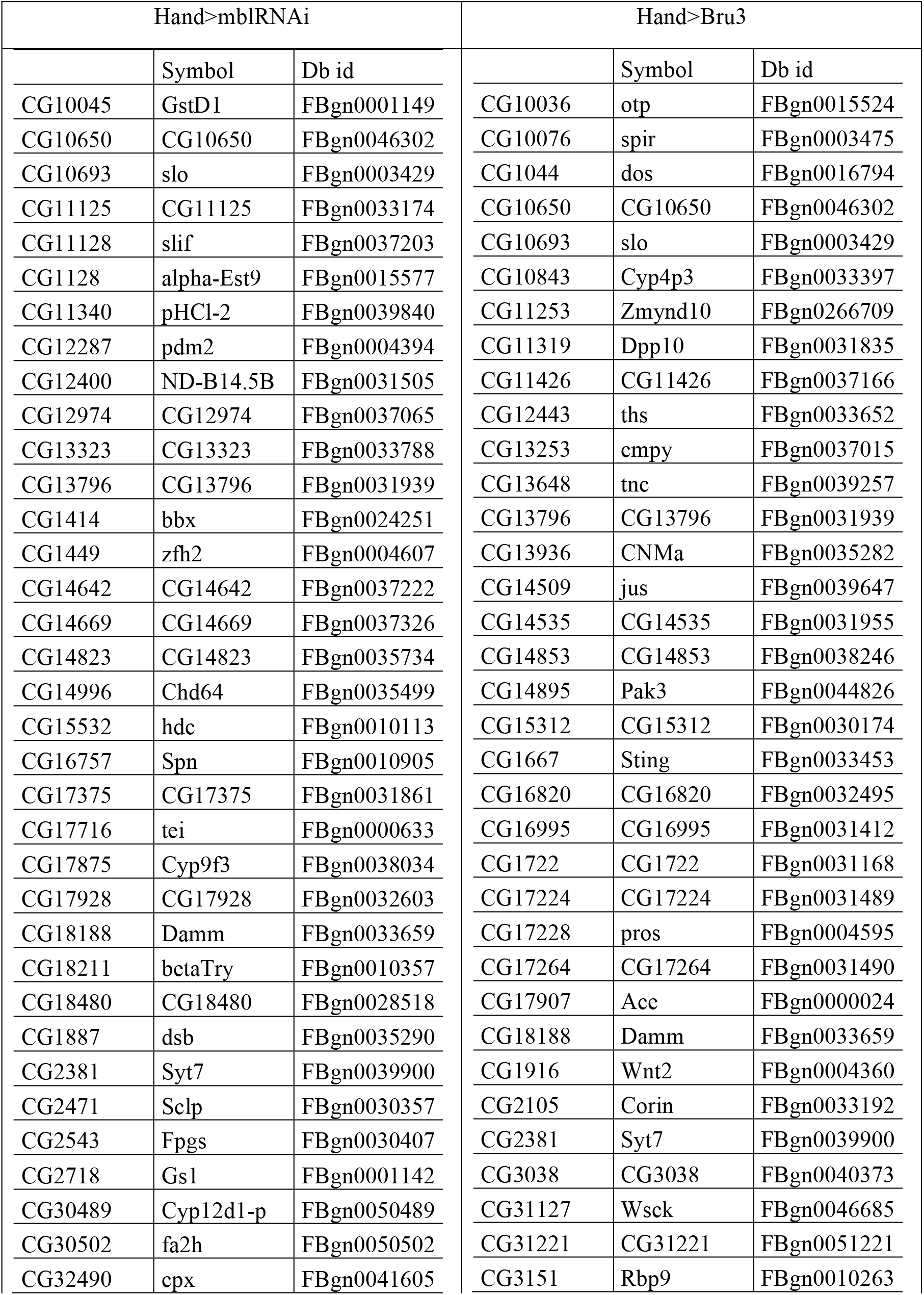
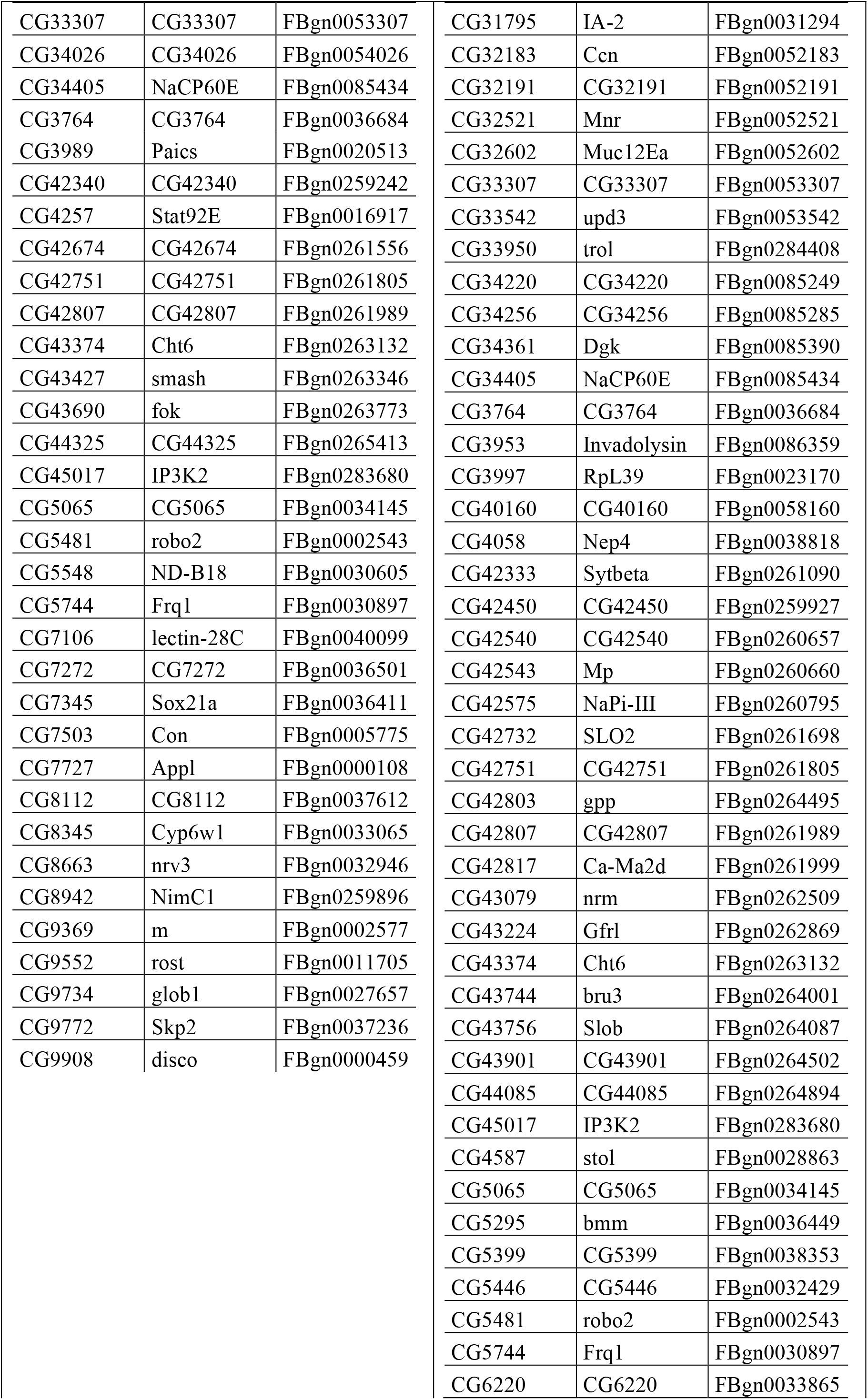
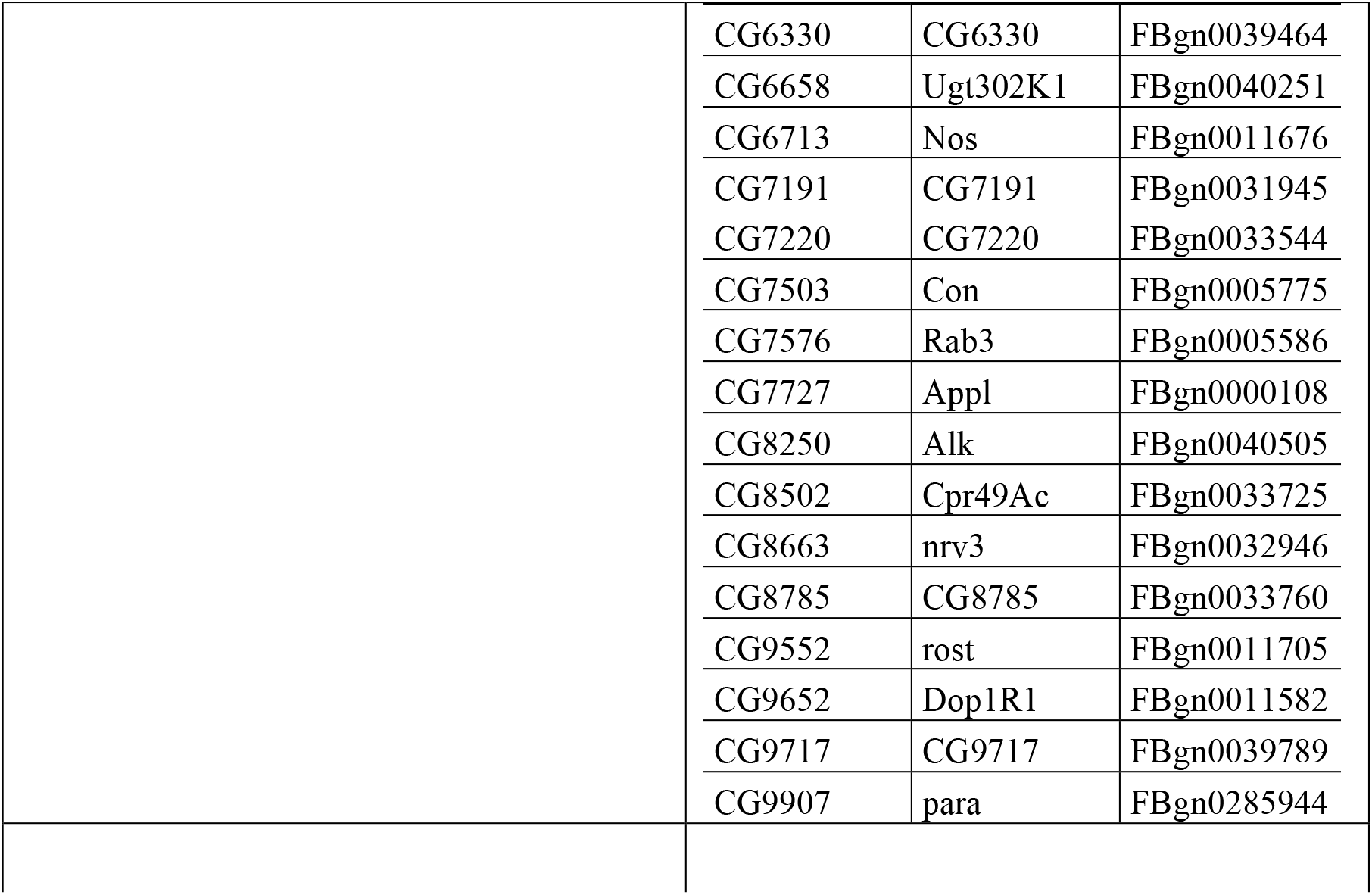
List of *in silico* identified dmiR-1 targets up-regulated in DM1 contexts developing DCM.

**Figure 3.**
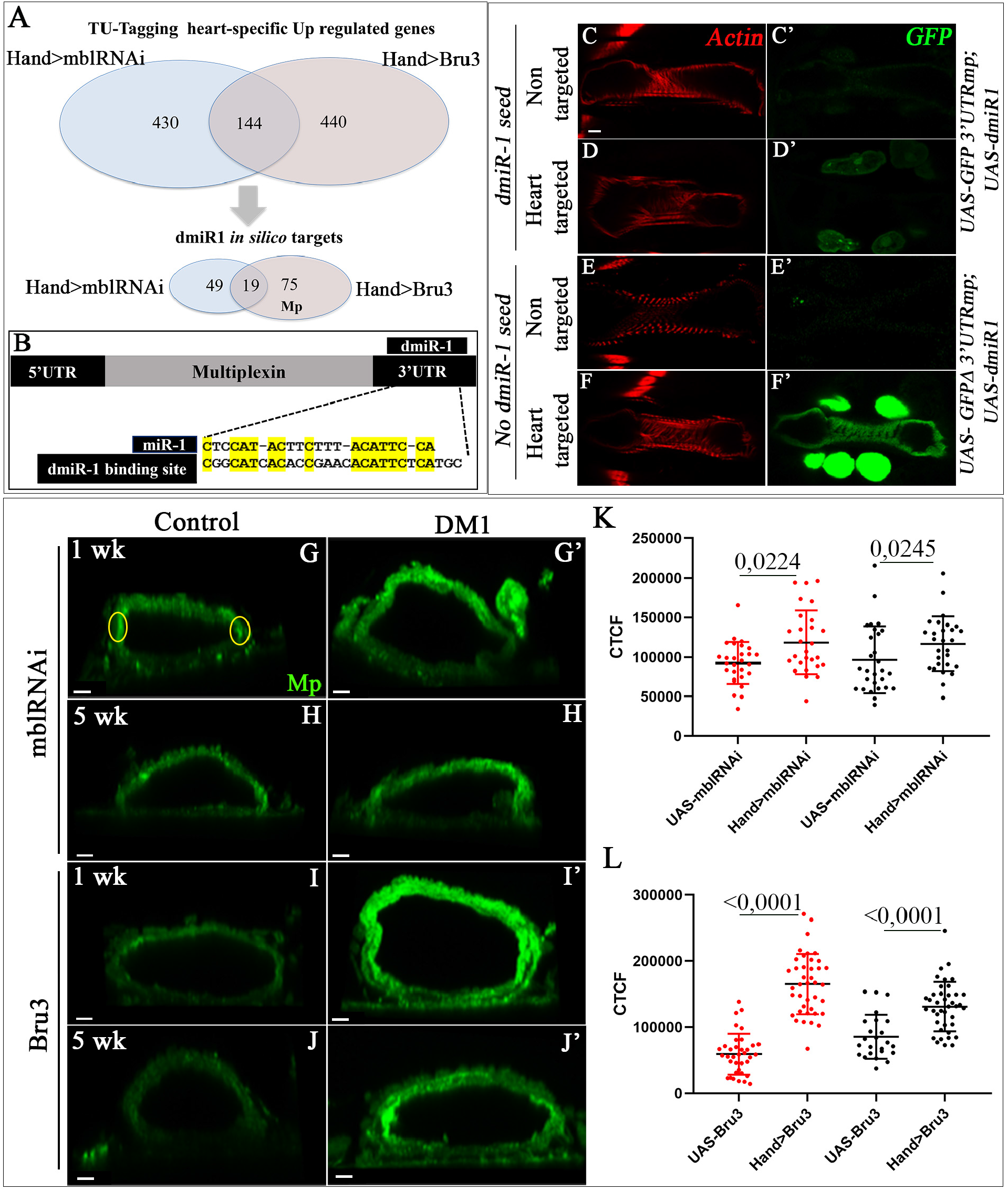
Multiplexin, a new cardiac *dmiR-1* target is up-regulated in DCM-developing DM1 flies. **(A)** Up-regulated genes identified by heart-specific transcriptomic approach (TU-tagging (Auxerre-Plantié et al., 2019) in DM1 contexts developing DCM (top Venn diagrams) underwent *in silico* screening for *miR-1* binding sites leading to identification of a set of potential *dmiR-1*-targets up-regulated in DM1 (lower Venn diagrams), of which *Multiplexin* with *miR-1* binding site in its 3’UTR region (*miR-1* seed site alignment) **(B)**. dmiR-1 binding site in *3’UTRMp* region is required to negatively regulate Mp expression *in vivo*. Adult hearts from transgenic GFP-sensor lines carrying *3’UTRMp* region with (*UAS_GFP 3’UTRMp)* or without *dmiR-1* seed site (*UAS_GFP Δ3’UTRMp*). These lines were combined with the *UAS-dmiR1* line to generate double transgenic lines and tested for GFP expression in non-targeted context **(C**,**E)** or in heart-targeted context **(D**,**F)** after crossing with cardiac Hand-Gal4 driver. When heart-targeted, GFP expression in *Hand>GFP 3’UTRMp; dmiR1* line (carrying *dmiR-1* seed site) **(D’)** is attenuated compared to *Hand>GFP Δ3’UTRMp; dmiR-1* line (lacking dmiR-1 seed site) **(F’)**. Scale bar = 40 µm. (**C)** Cross sections of adult hearts from one- and five-week-old controls (*UAS_mblRNAi, UAS_Bru3*) **(G**,**H**,**I**,**J)** and DM1 contexts (*Hand>mblRNAi, Hand>Bru3*) **(G’**,**H’**,**I’**,**J’)** labelled for Mp (green). Highlighted regions correspond to areas used for quantifications of the fluorescent signals **(G). (K**,**L)** Graphs of Mp signal quantification in cardioblasts from adult flies aged one and five weeks for controls (*UAS_mblRNAi, UAS_Bru3*) and DM1 contexts (*Hand>mblRNAi, Hand>Bru3*) using CTCF method. *n* = 9 hearts. Scale bar = 10 µm

*Mp*, a *Drosophila* ortholog of mammalian Collagen XVIII and Collagen XV, belongs to the family of multi-domain collagens. It is composed of an N-terminal thrombospondin-related domain, followed by multiple Collagen repeats, a Collagen trimerisation domain and a C-terminal endostatin domain (Meyer and Moussian, 2009). Mp also contains consensus glycosaminoglycan (GAG) attachment sites, and biochemical analysis of the protein extracted from embryonic tissues revealed the presence of chondroitin sulphate (CS) chains, making it more like human collagen XV than collagen XVIII in this respect. In the embryonic *Drosophila* heart, Mp is expressed in cardioblasts of the heart proper but not in aorta (Harpaz et al., 2013; Meyer and Moussian, 2009). The Mp was shown to be deposited in a polarised way along the heart lumen during cardiac tube formation (Meyer and Moussian, 2009) and involved in lumen shaping by enhancing Slit/Robo activity (Harpaz et al., 2013).

Here, we show that Mp is also expressed in the adult fly heart (Fig. S3B,B’,B’’). Consistent with the predicted location of an extracellular matrix protein, Mp was detected on the luminal and external surfaces of the cardiomyocytes ensuring cardiac contractions and was also present on the underlying adult heart ventral longitudinal muscles (VLM) (Fig. S3B,B’).

The predicted *dmiR-1-*binding site within the *Mp*-3’UTR region (Fig. 3B) is expected to negatively regulate *Mp* transcript level in the presence of *dmiR-1*. To assess the biological relevance of this binding site *in vivo*, we cloned the *Mp-3’UTR* fragment carrying the *dmiR-1* seed site downstream of the GFP coding sequence to generate the *UAS-GFP-Mp3’UTR* transgenic line. In parallel, the *UAS-GFP-ΔMp3’UTR* line was created in which the *dmiR-1* biding site was deleted from the 3’UTR *Mp* sequence. Both GFP sensor lines were then combined with the *UAS-dmiR-1* line to generate double transgenic lines *UAS-GFP-Mp3’UTR; UAS-dmiR-1* (Fig. 3C, D) and *UAS-GFP-ΔMp3’UTR*; *UAS-dmiR-1* (Fig. 3E, F), respectively. We found that expression of *dmiR-1* in *Hand>GFP-3’UTRMp* hearts significantly reduced the GFP signal in cardiac and pericardial cells (Fig. 3D’), suggesting that *dmiR-1* binds to the predicted seed site and negatively regulates *Mp* mRNA expression in the adult fly heart. The deletion of the *dmiR-1* binding site in *Hand>GFP-Δ3’UTRMp; dmiR1* heart prevented the repressive effects of *dmiR-1* (Fig. 3F’), demonstrating that the GFP silencing observed in *Hand>GFP-Mp3’UTR; dmiR1* hearts (Fig. 3D’) is *dmiR-1* dependent. This finding is also consistent with an increased Mp protein levels detected in hearts of the *Hand>miR-1-sponge* context mimicking *miR-1* attenuation and leading to DCM (Fig. S3J).

We then tested whether Mp protein level increased in DM1 contexts with DCM characterised by a reduced *dmiR-1*. We detected a significant increase in Mp protein level on the surface of cardiomyocytes in DCM-developing lines (*Hand>mblRNAi* and *Hand>Bru3*) in both young and aged flies (Fig. 3G,H,I,J,K,L). A similar increase in Mp expression was found in DM1 pericardial cells (Fig. S3D’,E’,F’,G’,H,I).

### Heart-targeted Mp overexpression leads to DCM in aged flies

Mp protein level increases in DCM-developing DM1 hearts and in the heart-specific *dmiR-1* attenuation context causing DCM. Moreover, the 3’-UTR region of *Mp* carries a *dmiR-1* binding site, making Mp an *in vivo dmiR1* target in the heart. All these observations prompted us to determine whether the increased cardiac Mp level could lead to DCM. To generate cardiac Mp gain-of-function we crossed *Hand-Gal4* with the *UAS-3HNC1 (UAS-Mp)* line (Meyer and Moussian, 2009). The *Hand>3HNC1* adult flies expressed a high Mp protein level in the heart (Fig. 4B,D). Already at age one week, the Mp gain-of-function flies displayed a larger heart tube diameter and a wider cardiac lumen than controls (*UAS-3HNC1)* (Fig. 4D). The diastolic and systolic diameters of hearts overexpressing Mp were also significantly increased relative to controls (Fig. 4E,F). However, cardiac contractility was not affected (Fig. 4G). In contrast, the five-week-old *Hand>3HNC1* flies, besides an increase in systolic and diastolic diameters (Fig. 4E,F,H,H’), showed significantly reduced fractional shortening and thus adversely affected cardiac contractility (Fig. 4G). These findings demonstrate that heart-specific increase in Mp protein level is detrimental to cardiac function, leading to DCM in aged flies.

**Figure 4.**
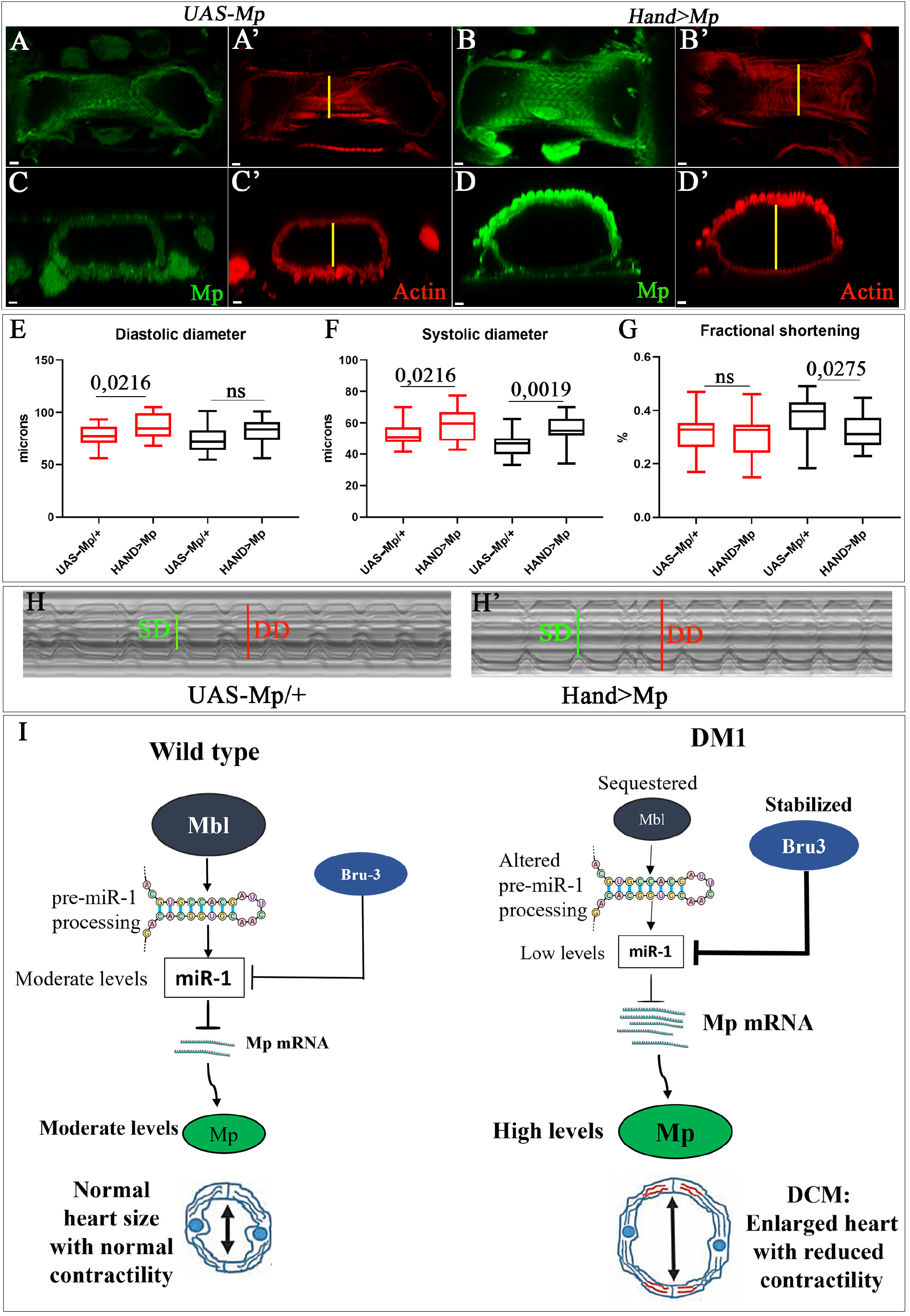
Heart-targeted Mp overexpression leads to DCM. **(A)** Adult heart aged one week **(A**,**B)** and its cross sections **(C**,**D)** labelled for Mp (green) and actin (red) for controls (*UAS_Mp*) and Mp overexpression context (*Hand>Mp*). Cardiac variables (diastolic **(E)** and systolic **(F)** diameters) and percent fractional shortening **(G)** for control (*UAS_Mp*) and Mp overexpression (*Hand>Mp)* conditions at ages one (red) and five weeks (black); *n* = 20 hearts. Scale bar = 5 µm. M-modes representing cardiac variables in five-week-old control **(H)** and Mp-overexpressing **(H’)** flies. **(I)** Scheme presenting cardiac role of *dmiR-1* and its target *Mp* in DCM-developing *Drosophila* DM1 models. In wild-type *Drosophila* heart Mbl promotes *pre-dmiR-1* processing. Bru-3 has potential antagonistic role in the destabilisation of *dmiR1*. As a result, in this context *dmiR-1* and its target Mp levels are moderate. In DM1, Mbl is sequestered and Bru-3 is stabilised, causing inefficient processing of *pre-dmiR1* and destabilisation of mature *dmiR-1*. As a result, in the DM1 context, the *dmiR-1* level is reduced while its target Mp level is high. This leads to an enlarged heart with adversely affected contractility.

To further analyse effects of Mp expression level on heart morphology and function, we tested the impact of cardiac-specific Mp knockdown at ages one and five weeks. We observed a reduced heart lumen in *Hand>MpRNAi* flies compared to controls and a significant decrease in diastolic and systolic diameters at ages one and five weeks, with no influence on cardiac contractility (Fig. S4).

Taken together, our findings demonstrate that Mp expression level plays a critical role in setting the size of the cardiac lumen and the systolic and diastolic fly heart variables, and its overexpression in the heart leads to DCM. We thus infer that the reduced *dmiR-1* leading to the up-regulation of its direct target Mp promotes development of DCM in the DM1 context (see scheme in Fig. 4I).

### Mp counterpart Col15A1 is up-regulated in cardiac samples of DM1 patients with DCM

Given the increased Mp level in DM1 fly models developing DCM, we examined whether expression of its human ortholog Col15A1 was also up-regulated in cardiac cells of DM1 patients. Of three DM1 cardiac tissue samples tested, two were from DM1 patients with DCM (Fig. 5A,B). At both RNA and protein levels, DM1 cardiac cells showed an increase in Col15A1 expression compared to cardiac cells from healthy donors, with differentially higher Col15A1 levels in DM1 patients with DCM compared to cardiac cells from DM1 patient without DCM (Fig. 5A,B). Thus, like Mp in *Drosophila* DM1 models, the up-regulation of Col15A1 in the heart correlates with DCM in DM1 patients.

**Figure 5.**
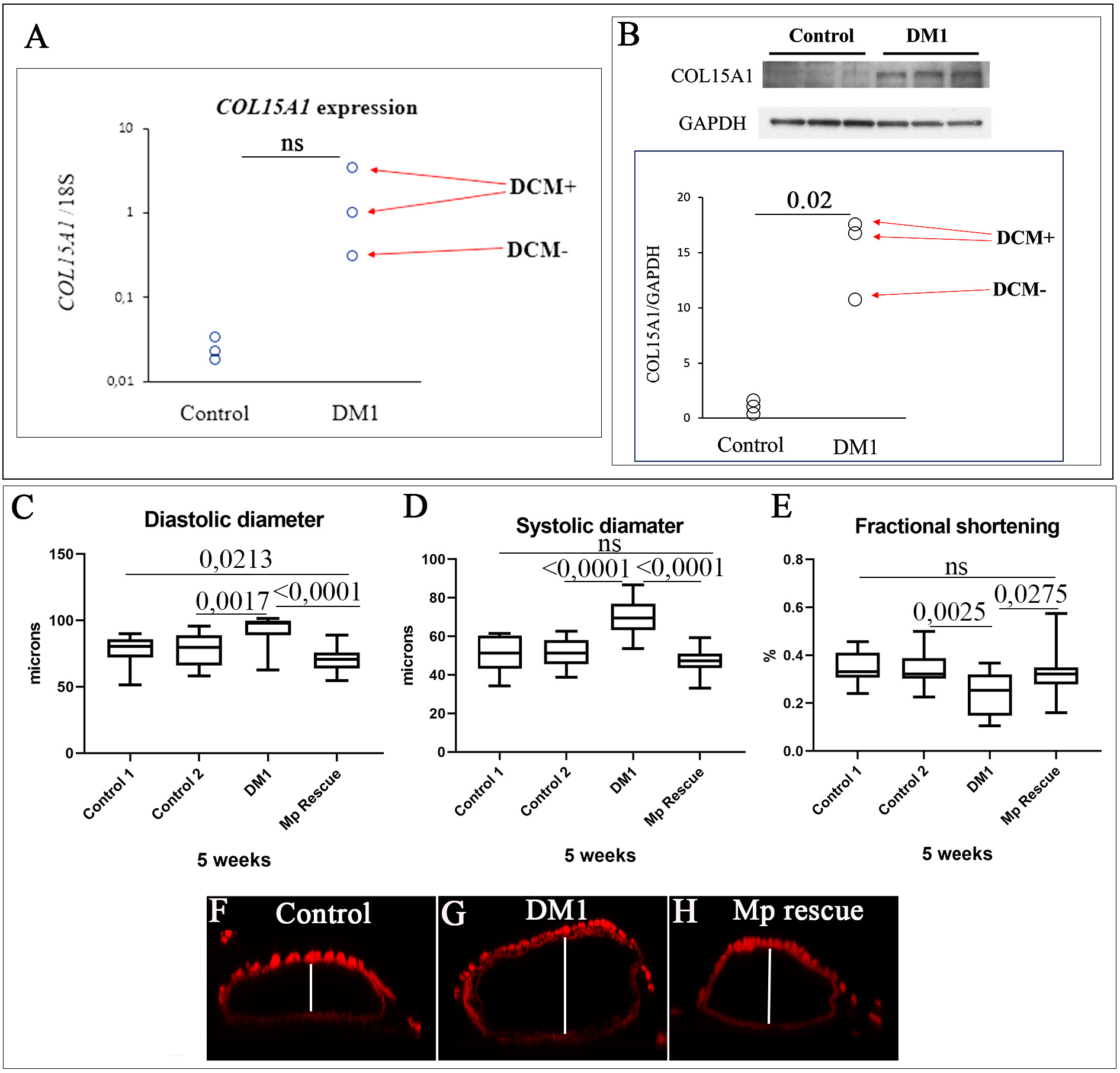
Increased cardiac expression of human Mp ortholog Col15A1 is associated with DCM in DM1 patients. **(A)** *Col15A1* transcript levels tested by RT-QPCR and **(B)** Col15A1 protein levels tested by Western blot in cardiac samples from healthy donors and from DM1 patients with and without DCM. Cardiac variables (diastolic **(C)** and systolic **(D)** diameters and percent fractional shortening **(E)**) assessed by SOHA approach for control 1 (*UAS-MpRNAi; UAS-Bru3)*, control 2 (*UAS-Bru3; UAS-UPRT*), Mp rescue (*Hand>MpRNAi; Bru3*) and DM1 context (*Hand>Bru3; UPRT*) at age five weeks. *n* = 20 hearts. Cross-sections of cardiac tubes 3D-reconstructed using Imaris software from five-week-old Control 1 (*UAS-MpRNAi; UAS-Bru3*) **(F)**, DM1 (*Hand>Bru3; UPRT*) **(G) and** Mp rescue (*Hand>MpRNAi; Bru3*) **(H)** flies labeled with actin. Scale barre = 20 µm

### Heart-specific attenuation of Mp ameliorates DM1-associated DCM phenotype

To determine whether the increased Mp/Col15A1 levels could contribute to the DCM phenotype in DM1, we applied our DM1 fly models and performed genetic rescue experiments by attenuating Mp expression in the DCM-developing *Hand>Bru3* context. In young one-week-old flies, heart-specific Mp attenuation had no effect on diastolic or systolic diameters, nor on contractility of *Hand>Bru3* hearts (Fig. S5). However, in five-week-old *Hand>Bru3; MpRNAi* flies, the systolic and diastolic diameters were reduced (Fig. 5C,D), associated with a significant increase in the fractional shortening variable compared to the aged DM1 context (Fig. 5E). However, rescue remained partial compared to aged control flies (Fig. 5E). Heart-specific attenuation of Mp thus improves DCM phenotype in aged DM1 flies.

## Discussion

Myotonic dystrophy type 1 is the most common muscular dystrophy in adults. Cardiac repercussions including DCM are among the main causes of death in DM1 (Groh et al., 2008). However, the underlying mechanisms remain poorly understood, impeding the development of adapted treatments. As we previously demonstrated (Auxerre-Plantié et al., 2019; Souidi et al., 2018; Souidi and Jagla, 2021), *Drosophila* DM1 models recapitulate all the cardiac phenotypes observed in DM1 patients and so could help gain insight into gene deregulations underlying DM1-associated DCM.

### DM1 fly models show reduced *dmiR-1* in cardiac cells and develop DCM

In humans, DCM is characterized by left ventricular dilation and systolic dysfunction defined by a depressed ejection fraction. Similarly, in DCM-developing flies, the cardiac tube is enlarged and shows an increased diastolic and systolic diameter with reduced contractility. The loss of cardiac miRNAs and in particular *miR-1* has already been correlated to DCM and heart failure in mice (Rao Prakash et al., 2009; Wei et al., 2014) and observed in patients with end-stage DCM (Ikeda et al., 2007). It is well known that *miR-1* regulates genes involved in cardiac development and function including *Nkx2*.*5, SRF* and components of *WNT* and *FGF* signalling pathways (Kura et al., 2020) and that its level is reduced in the pathological context of DM1 (Rau et al., 2011). However, it was not known whether the low *miR-1* level caused DM1-associated DCM, nor what were the downstream *miR-1* targets. Here, we show that two heart-targeting *Drosophila* DM1 models, *Hand>mblRNAi* and *Hand>Bru3* mimicking sequestration of MBNL1 and stabilisation of CELF1, respectively, developed DCM and showed a reduced expression of *dmiR-1* in cardiac cells. Because overexpression of CELF1 (Koshelev et al., 2010) and loss of MBNL1 (Lee et al., 2013) also result in DCM in mice, *Drosophila* appears well-suited to assessing the impact of reduced *miR-1* in DM1-associated DCM. One mechanism explaining why *miR-1* levels fall in the DM1 context is the sequestration of MBNL1, which can no longer play its physiological role in *pre-miR-1* processing into mature *miR-1* (Rau et al., 2011). Here, we observed reduced *dmiR-1* also upon cardiac overexpression of CELF1 ortholog Bru3. How CELF1/Bru3 impinges on *miR-1* levels is not fully understood, but the fact that CELF1 could bind UG-rich miRNAs and mediate their degradation by recruiting poly(A)-specific ribonuclease (PARN) (Katoh et al., 2015) could explain reduced *dmiR-1* levels in the *Hand>Bru3* context. Given that both *Drosophila* DM1 models developing DCM showed markedly reduced *dmiR-1* in cardiac cells, we sought to determine whether heart-targeted attenuation of *dmiR-1* was sufficient to induce DCM: *dmiR-1* knockdown in the heart mimics DM1-associated DCM.

### Col 15A1 ortholog Mp is a new *miR-1* target involved in DM1-associated DCM

To identify candidate *dmiR-1* target genes involved in DCM we performed *in silico* screening for *dmiR-1* seed sites in the 3’UTR regions of genes up-regulated in cardiac cells from DM1 models (Auxerre-Plantié et al., 2019) developing DCM (see Table 1). Among 1189 3’UTR sequences tested we found that 162 bore potential *dmiR-1* seed sites, of which the 3’UTR of *Multiplexin* (*Mp). Mp* codes for extracellular matrix protein belonging to a conserved collagen XV/XVIII family. We top-ranked *Mp* because of its known role in setting the size of the cardiac lumen (Harpaz et al., 2013). The *Mp-/-* embryos were found to present a narrower lumen with reduced contractility of the heart tube (Harpaz et al., 2013), whereas the mouse mutants of *Mp* ortholog, *Col15A1*, showed age-related muscular and cardiac deteriorations linked to a degraded organisation of the collagen matrix (Eklund et al., 2001; Rasi et al., 2010). This prompted us to examine how Mp was expressed in the adult fly heart and what the effect of its overexpression was. Using Mp specific antibody we detected Mp on the surface of the cardiac cells and found that it accumulated to a high level in both *Hand>mblRNAi* and *Hand>Bru3* DM1 lines. We also tested whether the *in silico* identified *dmiR-1* seed site was required for the regulation of Mp expression and confirmed that *Mp* is a direct *in vivo* target of *dmiR-1* in cardiac cells. Consistent with its role downstream of *dmiR-1*, Mp overexpression in the heart mimicked the *dmiR-1* knockdown phenotype, leading to a significantly enlarged heart with reduced contractility. Moreover, genetic rescue experiments with heart-specific attenuation of Mp expression in the *Hand>Bru3* DM1 context reduced heart dilation and ameliorated DCM phenotype in aged flies, thus demonstrating that increased Mp levels contribute to DCM observed in DM1 flies. Previous reports (Louzao-Martinez et al., 2016; Gil-Cayuela et al., 2016) revealed increased expression levels of different collagens associated with DCM in both animal models and patients. Here, we report evidence that Col 15A1 is specifically up-regulated at both transcript and protein levels in cardiac samples from DM1 patients and in particular in those with DCM. Altogether, the observations that Col15A1 expression level is abnormally elevated in DCM-developing DM1 patients and that attenuation of its *Drosophila* ortholog Mp could ameliorate the DCM phenotype suggest that Col15A1 could be a novel therapeutic target in DM1.

### DCM, a complex cardiac condition with a poor prognosis in DM1

A large number of genes have so far been implicated in DCM, attesting to the complex molecular origin of this cardiac condition. For example, in *Drosophila*, DCM was observed in mutants of genes encoding contractile and structural muscle proteins such as Troponin I (TpnI), Tropomyosin 2 (Tm2), δ-sarcoglycan and Dystrophin but also associated with deregulations of EGF, Notch, Cdc42 and CCR4-Not signalling pathway components (reviewed in Wolf, 2012). In humans, DCM-causing mutations were also identified in a large number of genes including those encoding cytoskeletal proteins such as FLNC, nuclear membrane protein LMNA or involved in sarcomere stability (Titin, TNNT2, MYH7, TPM1) but also RNA binding protein RBM20 (McNally and Mestroni, 2017).

Here, we focused on DCM associated with DM1. A previous study on a mouse model overexpressing CELF1 and developing DCM (Wang et al., 2015), identified down-regulation of *Transcription factor A mitochondrial* (*Tfam*), *Apelin* (*Apln*) and *Long-chain-fatty-acid-CoA ligase 1* (*Acsl1*) as potentially associated with DCM. It was suggested that CELF1 might regulate their mRNA stability by binding to their 3′UTR regions and causing destabilisation and degradation of their transcripts (Chang Kuei-Ting et al., 2017). In this DCM-developing mouse DM1 model, Col15a transcripts were elevated (Wang et al., 2015), but the role of Col15a in DCM was not analysed. Here, using *Drosophila* DM1 models with a DCM phenotype, we identified up-regulation of Col15A1 ortholog Mp as a molecular determinant of DM1-associated DCM. We also link reduced *miR-1* levels in DCM-developing DM1 cardiac cells to the up-regulation of Mp, establishing that *Mp* is an *in vivo* target of *miR-1*.

Importantly, our findings show that in DM1 patients, Collagen 15A1 is up-regulated in the hearts of patients with DCM. In DM1 patients, the DCM phenotype appears several years after onset and is less common than the conduction system defects and arrhythmias (Lin et al., 1989; Nguyen et al., 1988). However, it is frequently associated with poor prognosis and indication for heart transplant (Papa et al., 2018).

In summary, we report evidence for the importance of *miR-1*-dependent gene deregulations in DM1 and identify *Mp* as a new *miR-1* target involved specifically in DM1-associated DCM. We also demonstrate that Mbl depletion and Bru3 up-regulation in the heart have overlapping impacts on DM1 pathogenesis, both leading to reduced *miR-1*, up-regulation of Mp, and so to DCM (see scheme in Fig. 4C).

Our conclusion is that in a physiological context, Mp level is moderately triggered by Mbl-dependent regulation of *dmiR-1* processing and Bru-3-dependent regulation of *dmiR-1* stability. However, in the DM1 context, Mbl is sequestered in nuclear foci while Bru-3 levels increase, leading to a reduced *dmiR-1* and the up-regulation of its target gene *Mp*. Considering the known role of Mp as a positive regulator of cardiac lumen size (Harpaz et al., 2013) we would expect Mp accumulation in the adult heart also to promote heart tube enlargement, leading to the DCM phenotype. Whether like in embryos this Mp function involves the Slit/Robo signalling pathway remains to be investigated, but the finding that Robo2 is among identified *miR-1* targets up-regulated in DCM-developing DM1 flies (Table 1) supports this possibility. Finally, the fact that Mp ortholog Col15A1 is highly elevated in cardiac samples from DM1 patients with DCM and that reducing Mp ameliorates DCM phenotype in DM1 fly models suggests that Mp/Col15A1 could be an attractive diagnostic and/or therapeutic target for DM1-associated DCM.

## MATERIALS AND METHODS

### *Drosophila* stocks

All *D. melanogaster* stocks were grown and crossed at 25° C on standard fly food. In this study, we used three *Drosophila* DM1 models: *UAS-960CTG* (Picchio et al., 2013), *UAS-MblRNAi* (28732, VDRC Vienna, Austria) and *UAS-Bru3* (Harvard: d09837). For *dmiR-1* loss and gain of function we used *UAS-sponge dmiR-1* (Fulga et al., 2015) and *UAS-dmiR-1* (41125, Bloomington, USA) respectively. For functional analyses of Mp, we used *UAS-MpRNAi-TRIP* (52981, Bloomington, USA) and *UAS-3HNC1* (Meyer and Moussian, 2009). For testing the rescue of dilated cardiomyopathy observed in *Hand>Bru3* line, we have recombined *UAS-Bru3* with *UAS-MpTRIP* line to generate *UAS-MpTRIP; Bru3* line. The *UAS-Bru3* line was also recombined with the *UAS-UPRT* (UAS-HA-UPRT Bloomington 27604) line to generate *UAS-Bru3; UPRT* as a negative control for this rescue experiment.

All inducible lines were crossed with the driver line *Hand-Gal4* (provided by Laurent Perrin, TAGC, Marseille, France) to induce the expression of transgenes specifically in the fly heart (cardioblasts and pericardial cells). Control lines were generated by crossing the above-cited lines with *w*^*1118*^ line. *w*^*1118*^ is a mutant strain with a recessive white-eye phenotype.

### Heartbeat analyses of adult *Drosophila* hearts

Physiological analyses of adult *Drosophila* hearts were performed on one and five weeks old female flies using the SOHA approach protocol of Ocorr (Ocorr et al., 2009). For each genotype, about 20 flies were analyzed. First, we proceeded to dissection and preparation of semi-dissected hearts: adult *Drosophila* flies were anesthetized with Fly Nap for 5 minutes, then placed, dorsal side down, into a petri dish coated with a thin layer of petroleum jelly. The head, the ventral thorax and the ventral abdominal cuticle were removed in order to expose the abdominal contents. The ovaries and other organs as well as fats were then removed in order to expose heart tube attached to the dorsal cuticle. Dissection was performed in an oxygenated, artificial hemolymph (AH) solution composed of 108mM Na^+^, 5mM KCl, 2mM CaCl_2_, 8mM MgCl_2_, 1mM NaH_2_PO_4_, 4mM NaHCO_3_, 10mM sucrose, 5mM trehalose, 5mM HEPES (pH 7.1, all reagents from Sigma Chemicals). The solution was oxygenated by air-bubbling for 15 to 20 min. The beating hearts were filmed by a digital camera on 30 seconds movie with speed of 150 frames /second (Digital camera C9300, Hamamatsu, McBain Instruments, Chatsworth, CA) using microscope Zeiss (Axiophot, Zeiss) using immersion objective 10X. The heartbeats were recorded in A3 and A4 segments. The SOHA program based on Matlab R2009b software has been used for film analysis: The cardiac tube membrane during maximum diastole (relaxation) and maximum systole (contraction) were defined manually. One pair of marks identified the diastolic diameter, and one pair identified the systolic diameter. From this vertical row of pixels, a M-mode was generated to analyze the contraction and relaxation intervals: diastolic (DD), and systolic (SD) diameters, heart period (HP), and systolic (SI) and diastolic (DI) intervals. The diastolic and systolic diameters was used to calculate the Fractional shortening (FS) using the formula: (((Diastolic diameter - Systolic diameter)/Diastolic diameter) x 100) (Ocorr et al., 2009).

### Immuno-fluorescence staining of *Drosophila* heart

The adult hearts from one and five weeks old females were dissected as described previously, then fixed with formaldehyde 4%. Samples were incubated with primary rat anti-Mp antibodies (1/100) (Harpaz et al., 2013) or goat anti-GFP (1/500) (Abcam ab5450) overnight at 4 °C followed by 3 washes with PBS-Tween 0.1%, of 10 min each, then secondary antibodies anti-rat Alexa-CY3 (1/150) or with anti-goat Alexa-488 (1/150) (Jackson ImmunoResearch), respectively, for two hours at room temperature, Rhodamine phalloidin (1/1000) (Thermo Fischer Scientific) was used to reveal actin filaments. The preparations were mounted in a the Vectashield with DAPI (Vector laboratories, Inc. Burlingame, CA). Immunofluorescence labeled preparations were analyzed using a confocal microscope Leica SP8.

### Quantification of Multiplexin immunolabeling

Fluorescent-labelled heart tissues were all processed and stained under the same conditions. The level of fluorescent signal was quantified using ImageJ software by CTCF (Corrected Total Cell Fluorescence) approach. CTCF is calculated according to the formula: Measured by the software Integrated Optical Density - (Area of select x mean fluorescence of background readings). For each heart, CTCF was measured in two regions from segment A3 and two regions from segment A4 with the same areas for the 4 regions. For each genotype 9 hearts were analyzed for cardioblasts with 4 measurements in regions for each, and 3 pericardial cells were analyzed from each heart.

#### Single molecule fluorescence in situ hybridisation (smFISH)

For detection of *dmiR-1*, flies one and five weeks old were dissected and fixed for 30 min with 4% paraformaldehyde; rinsed at PBS1X -Tween 0.1% three-time, 5 min each. Samples were dehydrated through a series of increasing ethanol concentrations, transferred sequentially to 25%, 50%, 75%, 100% ethanol baths for 10 min each. After removing ethanol tissues were rehydrated through a series of decreasing ethanol concentrations, transferred sequentially 50%, 25% ethanol baths for 10 min each, and then post fixed for 30 min with 4% paraformaldehyde. Then incubated with solution composed of 50% PBT and 50% hybridisation buffer (5 mL Formamide, 0.5 mL SSC 20X, 100 µL ssDNA, 20 µL tRNA 50 ng/ µL, 5 µL Heparin 100 ng/µL, 10 µL Tween) for 5 min. The samples were finally incubated with hybridisation solution for 1h30 min at 50°C. The hybridisation mix including specific DIG-labeled probes (1nM for U6 snRNA positive control ref 339459, 20nM for negative control scramble ref 339459 and 40 nM for DME-miR1-3p ID: 339111) were added in each tube and samples were incubated at 50°C overnight. To remove non-specific RNA hybridisation, samples were washed with ISH (5 mL Formamide, 0.5 mL SSC 20X, 100 µL ssDNA, 5 µL Heparin 100 ng/µL, 10 µL Tween) for 30 min each at 50°C then with PBT. Slides were blocked in western blockage reagent (3mL blockage reagent+ 12mL PBT) for 30 min, incubated with sheep anti-DIG POD antibodies (11633716001, Roche) for 2 hours, treated with the TSA Plus Cyanine 3 System 1% (PerkinElmer, USA) for 5 min, blocked in 20% NGS diluted in PBT for 30 min. Finally, samples were incubated with primary antibodies (anti-Actin 1/250) overnight at 4 °C and then with secondary antibodies coupled with Alexa448 (1/150). The preparations were mounted in a Vectashield with DAPI medium. Immunofluorescence labeling preparations have been then analyzed using a confocal microscope Leica SP8.

### *sm*FISH quantification

Stained *Drosophila* hearts were imaged in 3D to allow the quantification of RNA transcripts in the total volume of cardiac and pericardial cells. The transcripts hybridized with *sm*FISH probes appears on the images as luminous fluorescent spots. The 3D images were analyzed using Imaris (version 9.3.1) that allows the detection, the visualization of spatial distribution as well as the quantification of intensity for each spot detected. To detect the real RNA spots of interest we have adjusted the threshold of intensity detection according to scramble. To determine the lower threshold in pericardial cells, we have analyzed the spots detected in 27 pericardial cells in 9 heart preparations labelled by scramble. We adjusted the lower treshold above the mean intensity of scramble spots (30) and the upper threshold (255) by default. With this background subtracting parameter, Imaris software generated *dmiR1* spots and calculated the mean intensity for each spot detected as well as the average of the mean intensities for all *dmiR-1* spots. Similar approach has been applied for dmiR-1 smFISH quantification in cardiac cells.

### RNA extraction and RT-qPCR on adult fly heart samples

Total RNA was isolated from 20 adult hearts from one and five weeks old female flies, using Direct-zol™RNA Microprep (ref: R2060) from Zymo Research following the manufacturer’s instructions. RNA quality and quantity were respectively assessed using Agilent RNA Screen Tape Assay on 4200 TapeStation System (Agilent Technologies) and Qubit RNA HS assay kit on a Qubit 3.0 Fluorometer (Thermo Fischer Scientific). Then, 150 ng total RNA was reverse transcribed using SuperScript IV Reverse Transcriptase kit (Invitrogen) with random primer mix, in a 20 µL reaction. Quantitative PCR was performed in 4 replicates in final volume of 10 µL using Light SYBR Green PCR Master Mix (Roche, Applied Science) on a LightCycler 480 Real-Time PCR System (Roche, Applied Science).2 µL (3 ng) of cDNA were added to a SYBR Green Master Mix. We used Rp49 as a reference gene. The following pairs of primers were used: *Rp49*: forward GCTTCAAGGGACAGTATCTG and reverse AAACGCGGTTCTGCATGAG; *pre-dmiR-1*: forward TTCAGCCTTTGAGAGTTCCATG and reverse CGCCAGATTTCGCTCCATAC. The relative quantifications of transcripts were obtained with the ΔΔCt method. Finally, nonparametric Mann–Whitney tests were performed to compare control samples and samples of interest.

### Generating transgenic fly lines

The *dmiR-1* binding site in 3’UTR of *Mp* was predicted by sequence alignment of 3’UTR *Mp* sequence from flybase (http://flybase.org/) to *dmiR-1* sequence from miRbase (http://www.mirbase.org/) using Bowtie2 (Langmead and Salzberg, 2012). To validate *Mp* as a direct target of *dmiR-1* in *vivo*, we have generated double transgenic fly lines. For the generation of *UAS-GFP-3’UTRmp* line, approximately 470 base pairs surrounding the predicted *dmiR-1*-target site in 3’UTR of *Mp* were amplified directly from genomic DNA from the *w*^*1118*^ flies using the primers Mp-F1: ATAACTAGTTGAGCGGAAACGGAAGGAAGAAGAGGAG and Mp-R1: ATATCTAGATGTTGTGAATGATGACGTTAGG and a high-fidelity DNA Polymerase enzyme (Thermo Scientific Phusion High-Fidelity DNA Polymerase) kit. For the *UAS-GFP-Δ3’UTRmp* line, the same steps were proceeded with primers Mp-F2: ATAACTAGTTGATAAAACAAAACAAATCACAGCAC and Mp-R1: ATATCTAGATGTTGTGAATGATGACGTTAGG to amplify about 350 bp without predicted *dmiR-1-*target site. The *Spe*I and *Xba*I restriction sites were incorporated into primers and introduced by PCR. PCR products were purified using NucleoSpin Plasmid clean up kit after validation of inserts by electrophoresis on 1% agarose gel. After digestion of purified *3’UTR Mp* fragments and *pUASP-PL-Venus* vector by *Spe*I and *Xba*I enzymes, we performed ligation between *3’UTR Mp* fragments and the vector using the T4 DNA ligase kit (Invitrogen) according to the manufacturer’s instructions. Purified vectors were then microinjected by the Fly Facility platform to generated transgenic lines. Finally, *UAS-GFP-3’UTRMp* and *UAS-GFP-Δ 3’UTRMp* transgenic lines were combined with *UAS-miR-1* (41125, Bloomington, USA) to generate *UAS-GFP-3’UTRMp; UAS-miR-1* and *UAS-GFP-Δ3’UTRMp; UAS-miR-1* respectively and then crossed with the driver line *Hand -GAL4*.

### RNA extraction, RT-qPCR, and immunoblot on DM1 hearts

Human ventricular cardiac muscle tissues were obtained at autopsy from 3 DM1 patients and 3 normal controls following informed consent. All experimental protocols were approved by the Institutional Review Board at Osaka University and carried out in accordance with the approved guidelines. Total mRNA was extracted and first-strand complementary DNA synthesized as described previously (Nakamori et al., 2008). RT-qPCR was performed using TaqMan Gene Expression assays (Hs00266332_m1 and 4333760F, Applied Biosystems) on an ABI PRISM 7900HT Sequence Detection System (Applied Biosystems), as described previously (Nakamori et al., 2011). Level of *COL15A1* mRNA was normalized to *18S* rRNA. For protein analysis, cardiac muscle tissues were homogenized in a 10x volume of radioimmunoprecipitation assay buffer (25 mM Tris-HCl; pH 7.5; 150 mM NaCl; 1% NP-40; 1% sodium deoxycholate; and 0.1% sodium dodecyl sulfate) containing a protein inhibitor cocktail (Sigma-Aldrich). The homogenate was centrifuged for 10 minutes at 10,000g and the supernatant was collected. Equal amounts of protein (40 µg) were separated by sodium dodecyl sulfate–polyacrylamide gel electrophoresis and transferred onto Immobilon-P membranes (Millipore), as previously described (Nakamori et al., 2008). Blots were blocked for nonspecific protein binding with 5% (w/v) nonfat milk and then incubated with a 1:500-diluted antibody against COL15A1 (Thermo Fisher Scientific) or 1:3000-diluted antibody against GAPDH (glyceraldehyde 3-phosphate dehydrogenase) (Sigma-Aldrich). After repeated washings, the membranes were incubated with horseradish peroxidase-conjugated goat anti-rabbit IgG (Thermo Fisher Scientific). The ECL Prime Western Blotting Detection Reagent (Cytiva) and ChemiDoc Touch Imaging System (Bio-Rad) were used to detect the proteins.

### Statistics

Nonparametric Mann-Whitney tests were performed to compare control samples and samples of interest of *Drosophila* model and t-test was performed to compare controls to DM1 context from heart samples of DM1 patients. All statistical analyses were performed using GraphPad Prism (version 8.0.1) software. Results are reported with *P<0*.*05* considered statistically significant.

