## Supplemental figure legends and supplemental figures for "Deregulations of miR-1 and its target Multiplexin promote dilated cardiomyopathy associated with myotonic dystrophy type 1"

**Legends to Supplementary Figures**

**Figure S1. *Hand>960CTG* flies do not develop DCM.** Cardiac size analyzes (diastolic **(A)** and systolic **(B)** diameters) and percent fractional shortening **(C)** performed by SOHA approach for controls (*UAS_960CTG*) and mutant (*Hand>960CTG*) at 1 (red) and 5 (darck) weeks of age. *n*= 20 hearts. (**D,E)** RT-qPCR analysis for *pre-dmiR1* transcript in adult heart of 1 and 5 weeks of age for control (*Hand>LacZ*) and DM1 contexts (*Hand>mblRNAi*, *Hand>Bru3*). *n*= 4 biological replicates.

**Figure S2. DM1 flies show reduced levels of *dmiR-1* in cardiac cells.**  smFISH for scramble (5’ 3’ DIG) **(A’)** and U6 (5’DIG) **(B’)** detected in *w^1118^* cardiac tube. smFISH for *dmiR-1* detection in controls (*UAS_dmiR1*, *UAS_mblRNAi*, *UAS_Bru3*) **(C’,E’,G’)** and mutants (*Hand>dmiR-1*, *Hand>mblRNAi*, *Hand>Bru3*) **(D’,F’,H’)** cardiac tubes labelled with *dmiR-1* probs (red) and actin (geen) at 1 week of age. Representative spot views generated using Imaris from *in situ* hybridisations with miRCURY LNA probe for *dmiR-1* and used for quantification of *dmiR-1* levels. Spot views of *dmiR-1* in hearts of one-week-old control (*UAS_mblRNAi*) **(I)** and DM1 (*Hand>mblRNAi*) flies **(I’)** are shown. The zoom area corresponds to pericardial cells. **(J,K)** Scatter plot graph showing the signal intensity quantified in pericardial cells of one- (red) and five- (black) week-old flies for controls (*UAS_Bru3*, *UAS_mblRNAi*) and DM1 contexts (*Hand>Bru3*, *Hand>mblRNAi*). *n* = 27 pericardial cells. Scale bar = 40 µm

**Figure S3. Mp is expressed in the adult fly heart and up-regulated in pericardial cells of DCM-developing DM1 lines. (A)** Adult heart of *w^1118^* line labeled for Mp (green), actin (red) and DAPI (Blue). **(C)** The yellow arrows indicate ventral longitudinal muscles (VLM). **(C’)** The yellow arrows indicate circular fibers (cardioblats). (**A’’, B’’)** Cross section in *w^1118^* cardiac tube, obtained after 3D reconstruction with Imaris software, showing expression of Mp in the internal and the external membrane of cardiac tube. Adult heart of 1 and 5 weeks of age labeled for Mp (green) for controls (*UAS_mblRNAi*, *UAS_Bru3*) **(D,E,F,G)** and DM1 context (*Hand>mblRNAi*, *Hand>Bru3*) **(D’,E’,F’,G’)**. Circled region in **E’** correspond to examples of areas used for quantifications of the fluorescent signal in pericardial cells using CTCF method.  **(H,I)** Fluorescence signal intensity quantification for Mp expression in perdicardial cells in adult heart of 1 and 5 weeks of age for controls (*UAS_mblRNAi*, *UAS_Bru3*) and DM1 context (*Hand>mblRNAi*, *Hand>Bru3*) using CTCF method. Scale barre = 20µm. **(J)** Fluorescence signal intensity quantification for Mp expression in cardioblasts in adult heart of one week of age for control (*UAS_dmiR-1-sponge*) and mutant (*Hand>dmiR-1-sponge*) using CTCF method.

**Figure S4. Mp loss of function leads to reduced heart size.** Adult heart of 1 week of age labeled for Mp (green) and actin (red) for control (*UAS_Mp RNAi TRIP52981*) **(A,A’)** and mutant (*Hand>Mp RNAi TRIP52981*) **(B,B’)**. **(C)** Fluorescence signal intensity quantification for Mp expression in cardioblasts in adult heart of 1 week old for control (*UAS_Mp RNAi TRIP52981*) and mutant (*Hand>Mp RNAi TRIP52981*) using CTCF method. Cardiac size analyzes (diastolic (**D)** and systolic diameters **(E)**) and percent fractional shortening **(F)** performed by SOHA approach for control (*UAS_Mp RNAi TRIP52981*) and mutant (*Hand>Mp RNAi TRIP52981*) at 1 (red) and 5 (darck) weeks of age. *n* = 20 hearts. Scale barre = 20µm

**Figure S5. The inhibition of Mp has no effects of DCM in DM1 young heart.** Cardiac size analyzes (diastolic **(A)** and systolic **(B)** diameters) and percent fractional shortening **(C)** performed by SOHA approach for control 1 (*UAS_Mp RNAi TRIP52981; Bru3)* and control 2 (*UAS_Bru3; UPRT*) and Mp rescue (*Hand>Mp RNAiTRIP52981; Bru3*) and DM1(*Hand>Bru3; UPRT*) at 1 week of age. *n* = 20 hearts.

**Figure S1**

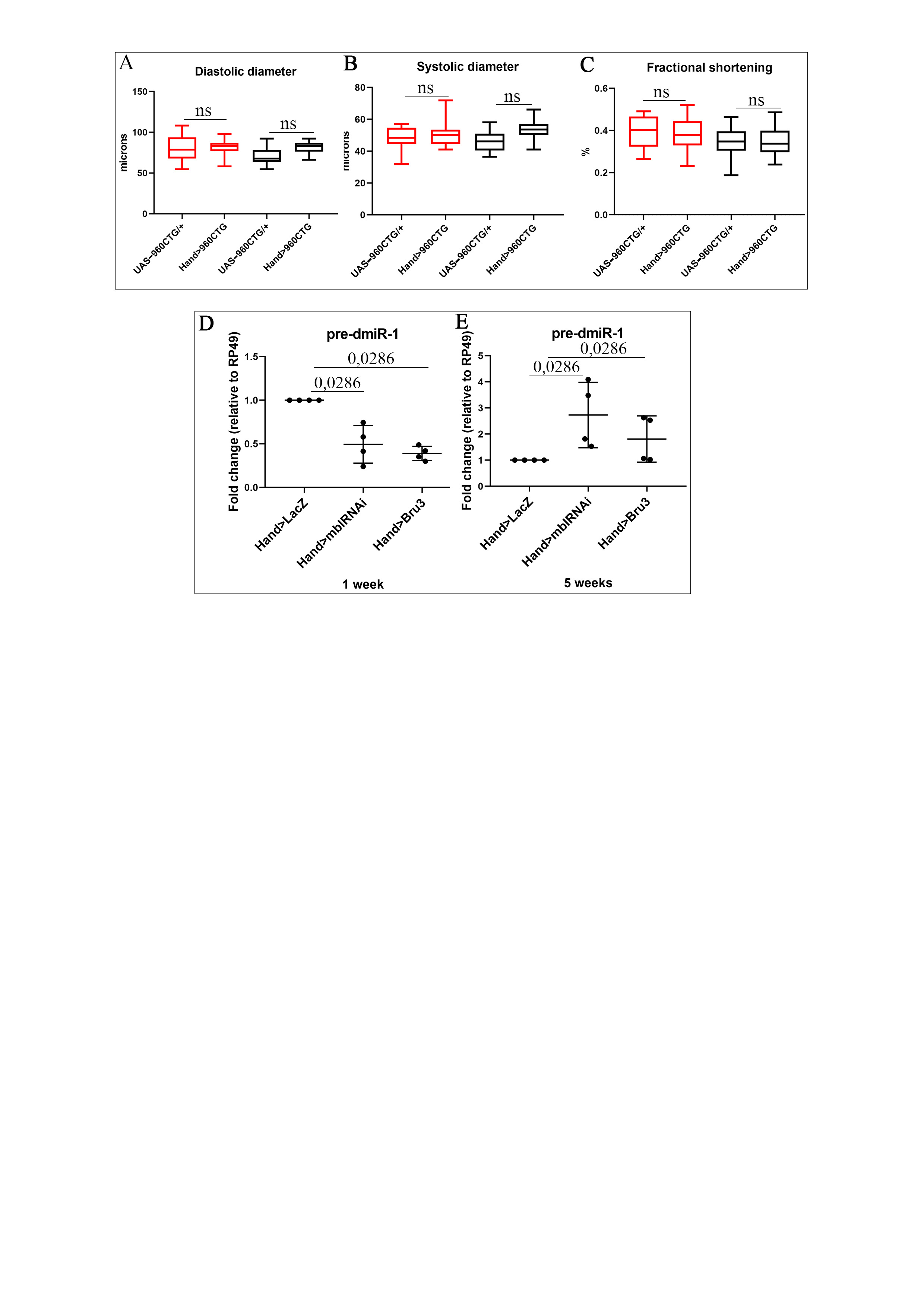

**Figure S2**

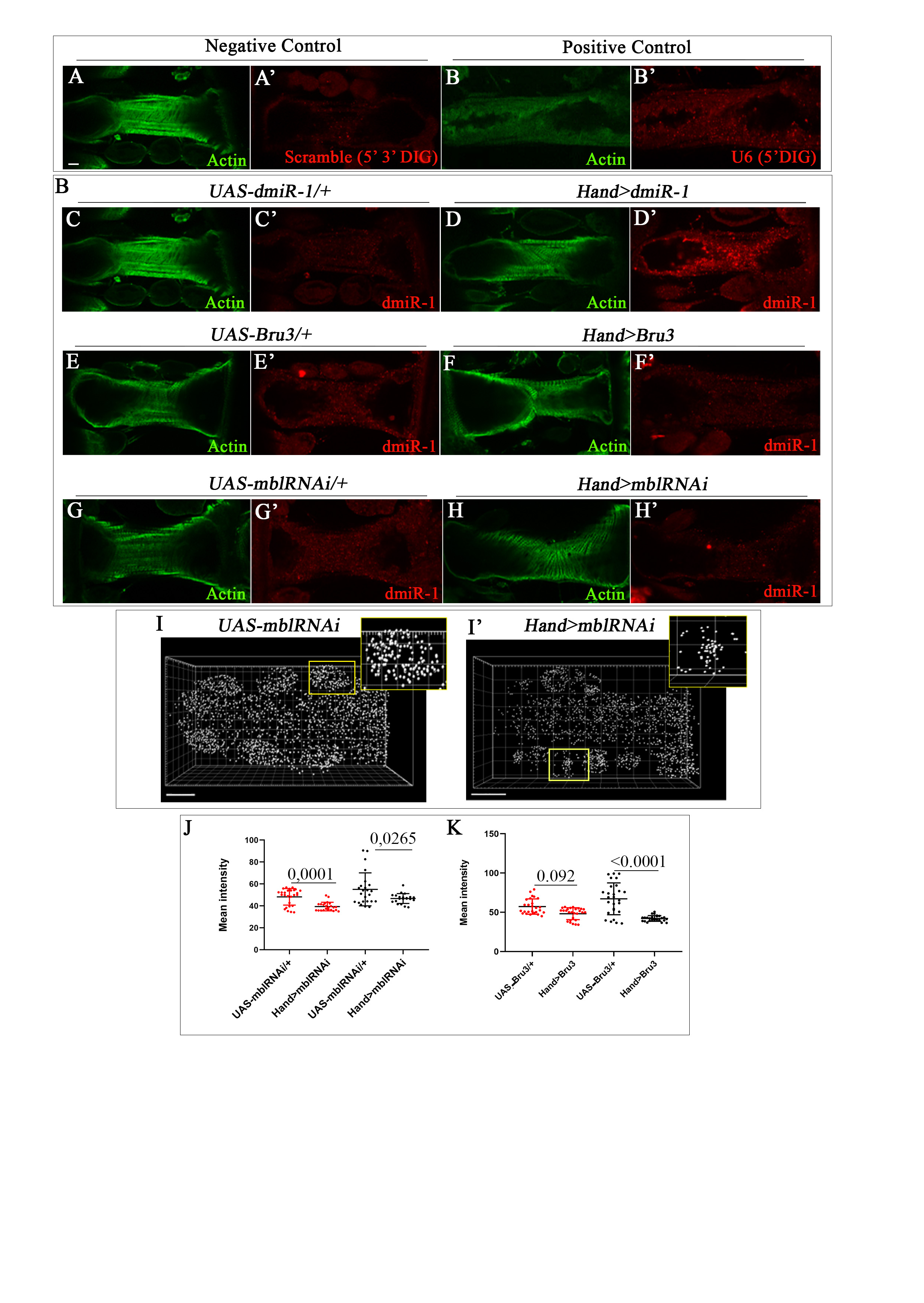

**Figure S3**

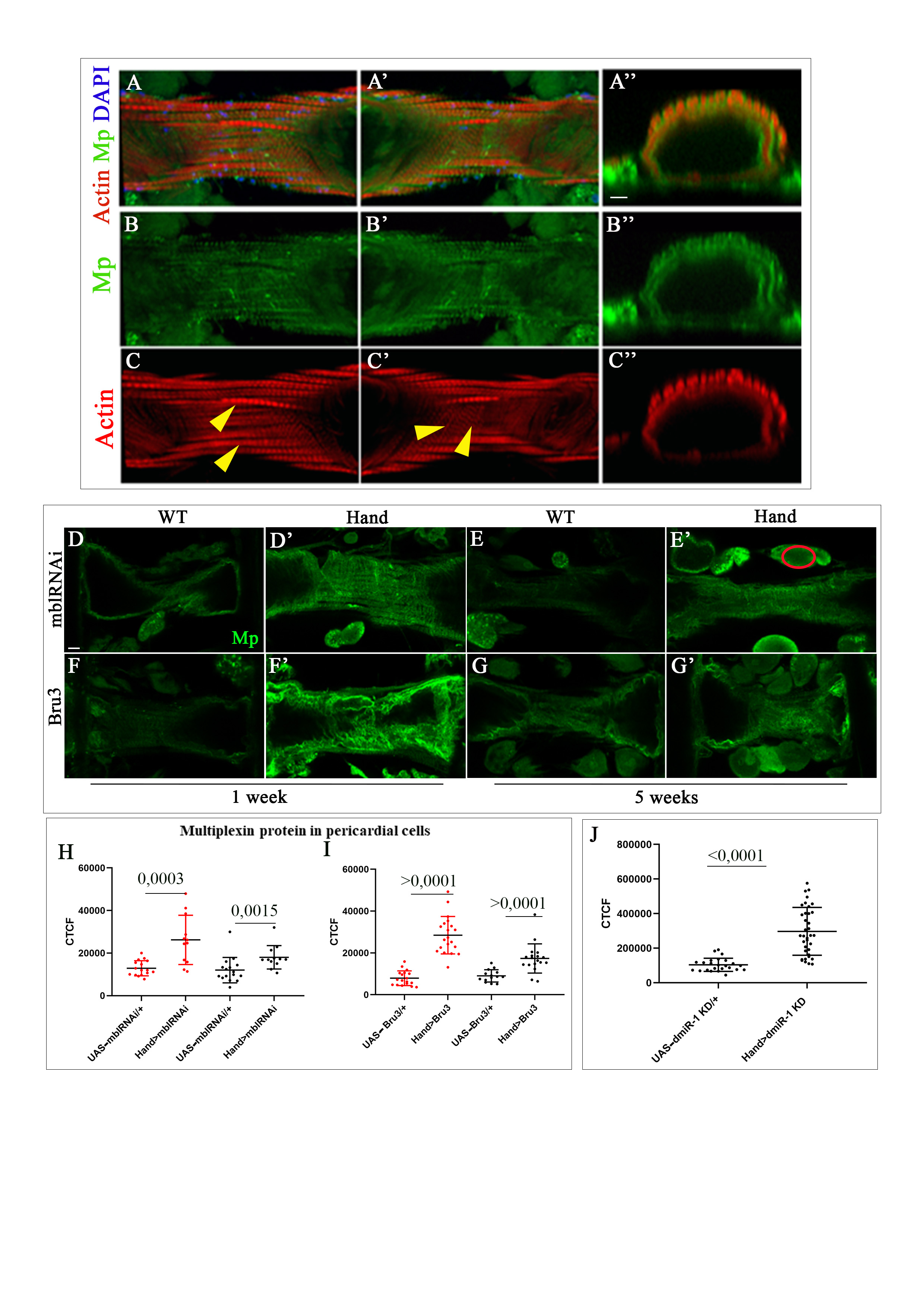

**Figure S4**

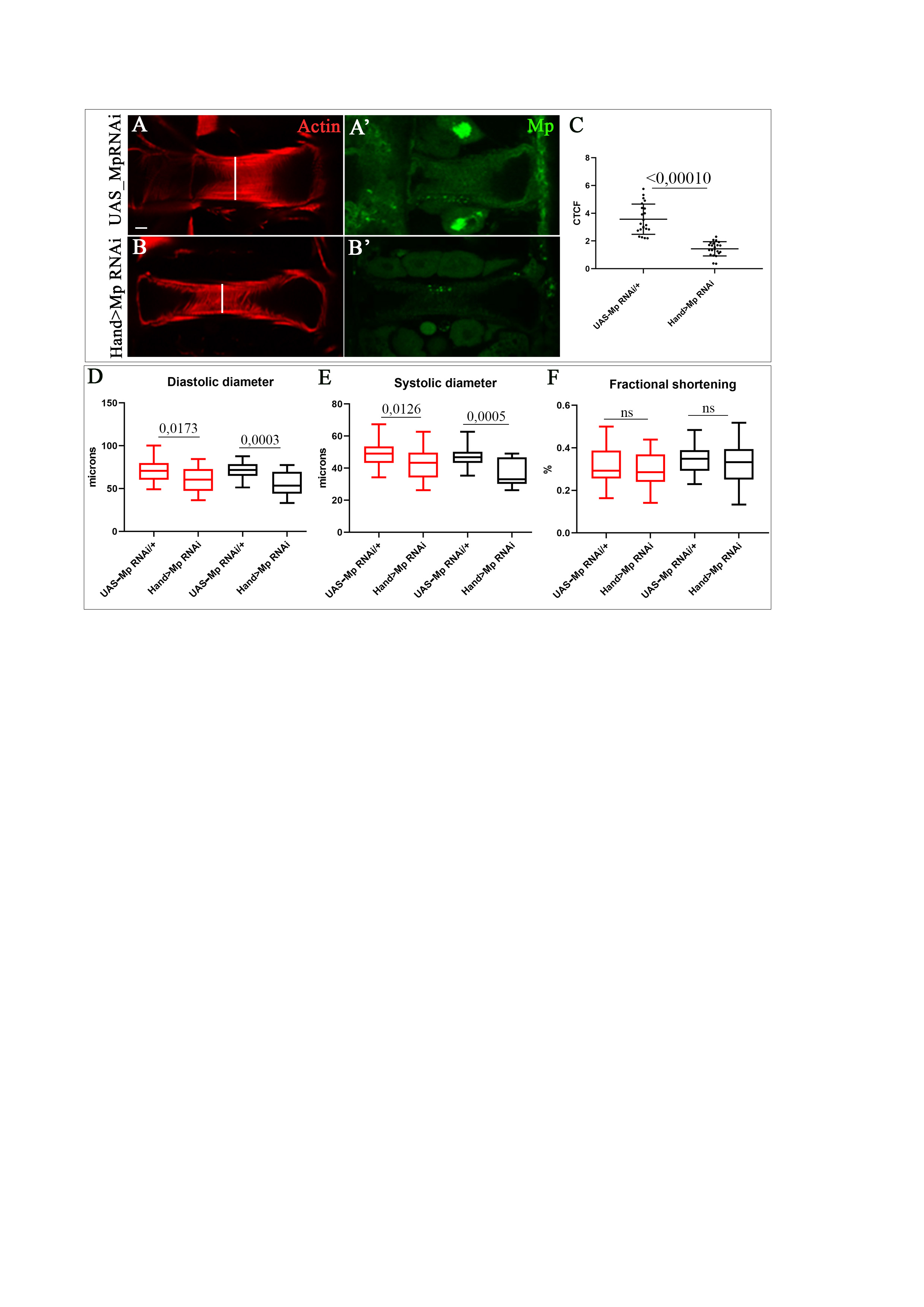

**Figure S5**

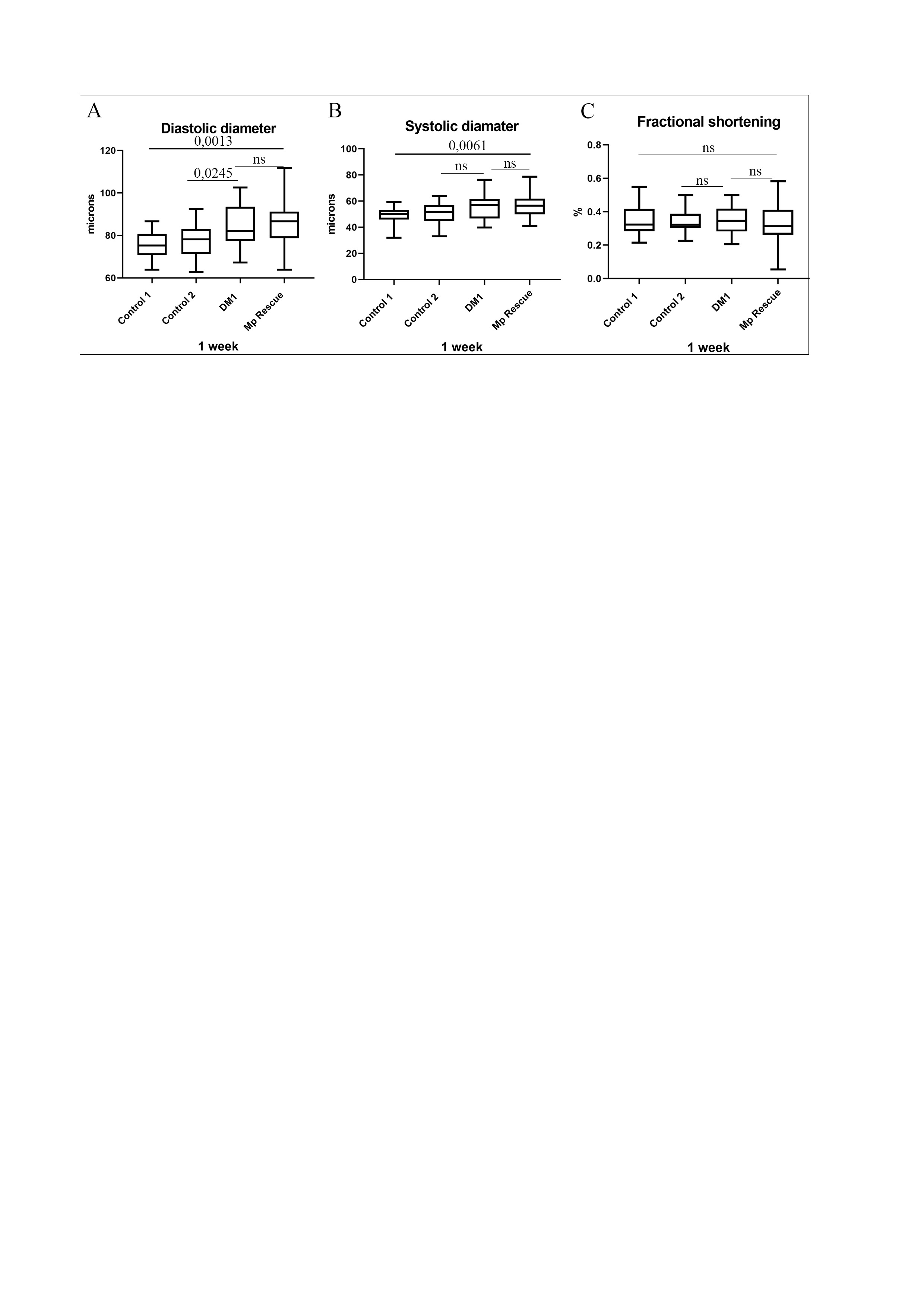
